# scCODA: A Bayesian model for compositional single-cell data analysis

**DOI:** 10.1101/2020.12.14.422688

**Authors:** M. Büttner, J. Ostner, CL. Müller, FJ. Theis, B. Schubert

## Abstract

Compositional changes of cell types are main drivers of biological processes. Their detection through single-cell experiments is difficult due to the compositionality of the data and low sample sizes. We introduce scCODA (https://github.com/theislab/scCODA), a Bayesian model addressing these issues enabling the study of complex cell type effects in disease, and other stimuli. scCODA demonstrated excellent detection performance and identified experimentally verified cell type changes that were missed in original analyses.

## Main

## Introduction

Recent advances in single-cell RNA-sequencing (scRNA-seq) allow large-scale quantitative transcriptional profiling of individual cells across a wide range of tissues, thus enabling the monitoring of transcriptional changes between conditions or developmental stages and the data-driven identification of distinct cell types.

Although being important drivers of biological processes such as in disease^1^, development^2^, aging^3^, and immunity^4^, shifts in cell type compositions are non-trivial to detect using scRNA-seq. Statistical tests need to account for multiple sources of technical and methodological limitations, including low number of experimental replications. The total number of cells per sample is restricted in most single-cell technologies, implying that cell type counts are proportional in nature. This, in turn, leads to a negative bias in cell type correlation estimation^5^ (**Fig. 1a**). For example, if only a specific cell type is depleted after perturbation, the relative frequency of others will rise. If taken at face value, this would lead to an inflation of differential cell types. Therefore, standard univariate statistical models that test compositional changes of each cell type *independently* may falsely deem certain population shifts as real effects, even though they were solely induced by the inherent negative correlations of the cell type proportions. Yet, common statistical approaches currently applied in compositional cell type analysis ignore this effect. For example, Haber et al.^6^ applied a univariate test based on Poisson regression, Hashimoto et al.^3^ a Wilcoxon rank-sum test, and Cao et al.^7^ proposed a method based on a generalized linear regression framework with a Poisson likelihood, all thus not addressing the issue of compositionality.

**Figure 1.**
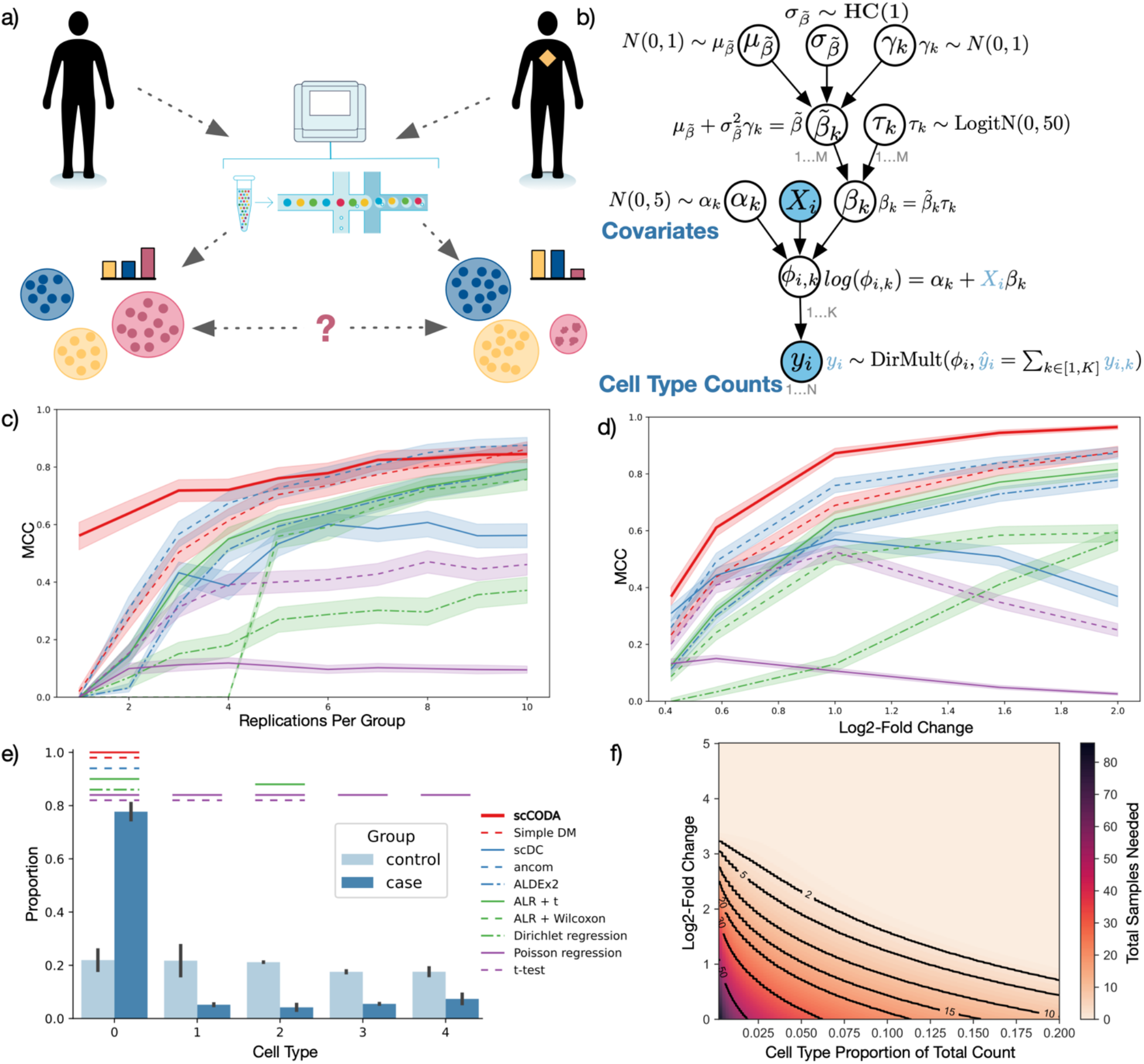
Compositional data analysis in single-cell RNA-sequencing data. **(a)** Single-cell analysis of control and disease states of a human tissue sample. Disease states reflect changes in the cell type composition. **(b)** The scCODA model structure with hyperparameters. Blue variables are observed. *DirMult* indicates a Dirichlet-Multinomial, *N* a Normal, *logitN* a Logit-Normal, and *HC* a Half-Cauchy distribution. **(c-d)** Benchmark of scCODA’s performance over all tested scenarios measured by Matthews’ correlation coefficient (MCC). Bayesian models (red), non-standard compositional models (blue), compositional tests/regression (green), non-compositional methods (purple). **(c)** Performance with increasing effect size shown as log2-Fold Change. **(d)** Performance with an increasing number of replicates. **(e)** Exemplary realization of the tested scenarios with high compositional log-fold change and low replicate number. Bars indicate statistically detected compositional changes between case and control for different methods. **(f)** Power analysis model to determine the number of replicates required to detect a significant change in cell type compositions dependent on the cell type abundance (x-axis) and the effect size shown as log2-Fold Change (y-axis).

To account for the inherent bias present in cell type compositions, we drew inspiration from methods for compositional analysis of microbiome data^8,9^ and propose a Bayesian approach for cell type composition differential abundance analysis to further address the low replicate issue. The **s**ingle-**c**ell **co**mpositional **d**ata **a**nalysis (scCODA) framework models cell type counts with a hierarchical Dirichlet-Multinomial distribution that accounts for the uncertainty in cell type proportions and the negative correlative bias via joint modeling of all measured cell type proportions instead of individual ones (**Fig. 1b**, **Online Methods - Model**). The model uses a Logit-normal spike-and-slab prior^10^ with a log-link function to estimate effects of binary (or continuous) covariates on cell type proportions in a parsimonious fashion. This implicitly assumes that only few cell types change upon perturbation. Since compositional analysis always requires a reference to be able to identify compositional changes^5^, scCODA currently relies on the manual specification of a reference cell type^11^. This implies that credible changes detected by scCODA have to be interpreted in relation to the selected reference.

## Results

### scCODA performs best in a benchmark of synthetic datasets

We first performed comprehensive benchmarks on synthetic data across a wide range of scenarios (**Online Methods - Simulation**) that focused on scCODA’s primary application: the behavior of a single binary covariate that models the effect of a perturbation of interest in the respective scRNA-seq experiment. To detect statistically credible changes in cell type compositions, we calculate the model inclusion probability for each covariate determined by the spike-and-slab prior **(Online Methods - Model description**). The benchmark scenarios were then used to identify the optimal inclusion probability threshold to deem a covariate statistically credibly affecting cell type composition. We found that the optimal inclusion threshold is roughly proportional to the number of cell types (**Supplementary Fig. 1a-e**). This reflects the necessity to have higher confidence in the identified effect with increasing number of cell types, mimicking the behavior of frequentistic multiple testing correction.

We next compared scCODA’s performance to state-of-the-art differential compositional testing schemes from the microbiome field as well as all non-compositional tests recently applied to single-cell data (**Fig. 1c-e**). In our synthetic benchmarks, we found scCODA to significantly outperform all approaches across a wide variety of effect sizes and experimental settings with an average Matthews correlation coefficient (MCC) of 0.75. Considering effect sizes, scCODA showed the highest MCC across all scenarios and consistent improvement with effect size strength (**Fig. 1c**). Other compositional (Bayesian) models such as a standard Dirichlet Multinomial (DM), ANCOM^12^, ALDEx2^13^, and additive log-ratio (ALR) transformed proportions combined with a t-test **(Online Methods - Model Comparison)** showed similar behavior, albeit with lower MCC. Considering the number of replicates per group, scCODA had a considerable edge over other methods in the common scenario with low number of replicates per group (seven or less) (**Fig. 1d**). ANCOM and the standard DM model achieved comparable performance with higher number of replicates, outcompeting ALDEx2 and ALR-transform models. Non-compositional models (purple lines in **Fig. 1c-e**) included more false-positives with increasing effect size (exemplified in **Fig. 1e**) and number of replicates per group, highlighting the detrimental effect of compositional data on these models.

### Power analysis to detect compositional changes

Since extensive replication of scRNA-seq experiments is still costly and hence rare, yet essential for studying compositional changes, we also investigated the sample size dependency of effect size and rarity of affected cell type on scCODA’s performance (**Fig. 1f**). We performed a power analysis fitting a linear model (R^2^=0.816±0.006, **Online Methods - Power Analysis**) on empirical logit transformed^14^ and scaled MCC values to infer the required sample size to reach an MCC of 0.8 for varying log-fold changes (**Fig. 1f, Supplementary Fig. 1f-g**). We estimated that a relative change of 1 (log2 scale) in abundant cell types (e.g., 1,000 out of 5,000 cells) can be determined with two samples, while the same relative change requires between 20 and 30 samples in a rare cell type (e.g., 125 out of 5,000 cells). Notably, large relative changes (log-fold changes of 3) in rare cell types could be detected from only two samples. While this implies that for many situations only two replicates are useful, we would advise to increase the number of samples when detection of compositional changes in rare cell types is relevant.

### scCODA identifies the FACS verified decrease of B-cells in supercentenarians

Next, we applied scCODA to a number of scRNA-seq data examples^1,3,4,6,15^ (**Fig. 2** and **Supplementary Fig. 2-5**). To confirm scCODA’s applicability on real data with known ground truth, we first considered a recent study of age-related changes in peripheral blood mononuclear cells (PBMCs)^3^, where cellular characteristics of supercentenarians (n=7) were compared against the ones of younger controls (n=5; **Fig. 2a**). The original study used a Wilcoxon rank-sum test and reported a significant decrease of B-cells in supercentenarians, which is known from literature^16^. Moreover, the result was validated by FACS measurements. scCODA also identified B-cell populations as the sole affected cell type using Erythroblasts as reference. The effect could be consistently detected with only three samples per condition (in 7/10 stratified random subsamples) while the Wilcoxon rank-sum test failed to detect the effect when removing even just one sample from each condition. This suggests that scRNA-seq data indeed comprise enough information to study compositional changes, and that scCODA can correctly identify the experimentally validated age-related decrease of B-cells even in low sample regimes.

**Figure 2.**
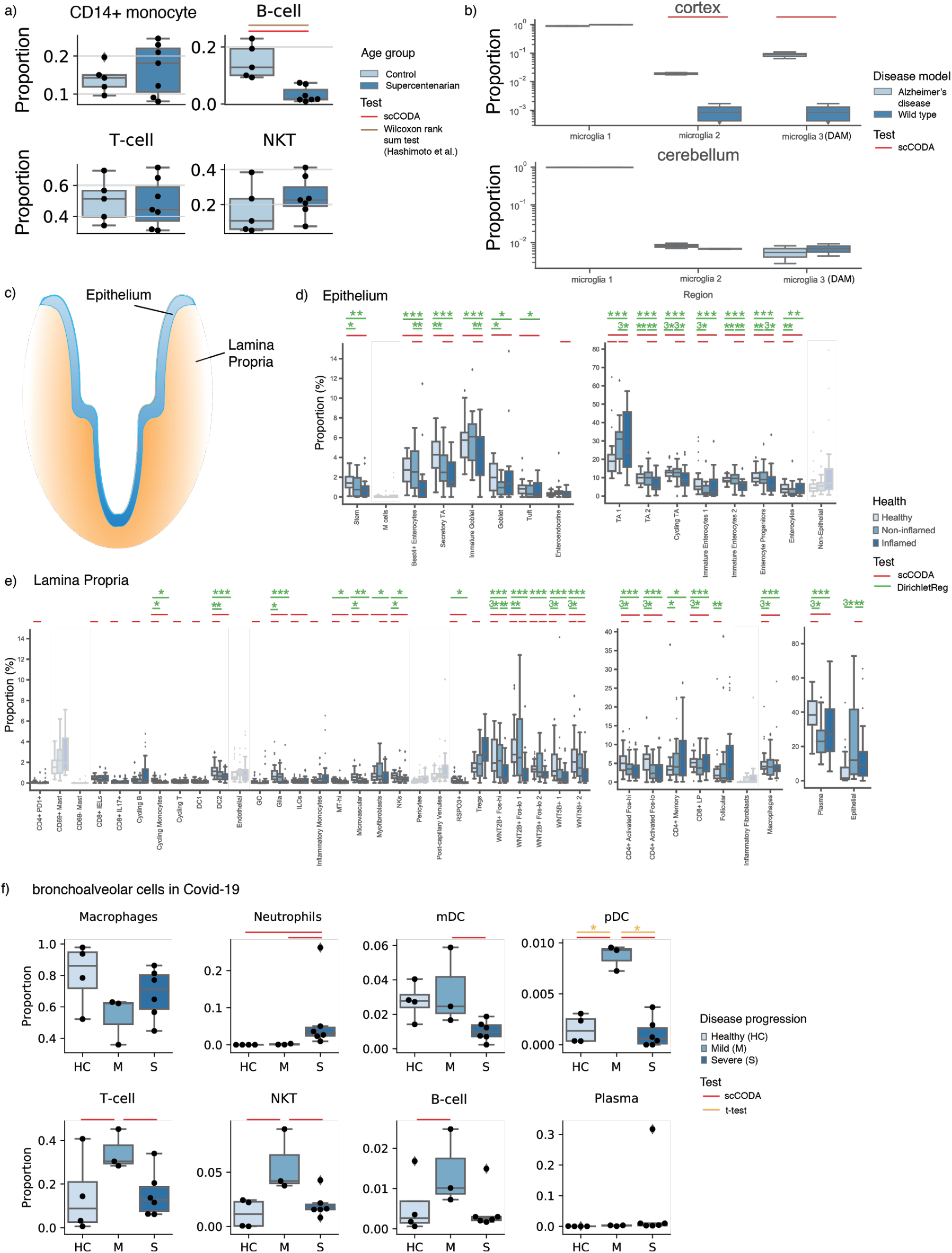
scCODA determines the compositional changes in a variety of examples. **(a)** Blood samples of supercentenarians (n=7) have significantly fewer B cells than younger individuals (control, n=5), reference was set to Erythroblasts, Hamiltonian Monte Carlo (HMC) chain length was set to 20,000 with a burn-in of 5,000. Credible and significant results are depicted as colored bars (Red: scCODA, brown: Wilcoxon rank-sum test^3^). Results are in accordance with FACS data^3^. **(b)** Microglia associated with Alzheimer’s disease (AD) are significantly more abundant in the cortex, but not in cerebellum^15^ (n=2 in AD and wild type mice, respectively), reference was set to microglia 1, HMC chain length was set to 20,000 with burn-in of 5,000. **(c-e)** Changes in epithelium and lamina propria in the human colon^1^ in ulcerative colitis (UC) (n=133 from 18 UC patients, 12 healthy donors). Credible and significant results are depicted as colored bars (Red: scCODA, green: Dirichlet Regression). Stars indicate the significance level (*: adjusted p<0.05, **: adjusted p<0.01, ****: adjusted p<0.001; Benjamini-Hochberg corrected). **(c)** Epithelium and Lamina propria are distinct tissues, which are studied separately. **(d)** Compositional changes from healthy to non-inflamed and inflamed biopsies of the intestinal epithelium, reference was set to Non-Epithelial cells, HMC chain length was set to 150,000 with burn-in of 10,000. **(e)** Compositional changes from healthy to non-inflamed and inflamed biopsies in the lamina propria, reference was set to six different cell types (see Methods), HMC chain length was set to 400,000 with burn-in of 10,000. **(f)** Compositional changes in bronchoalveolar cells in COVID-19 patients (n=4 healthy, n=3 mild, n=6 severe disease progression)^4^. Credible and significant results are depicted as colored bars (Red: scCODA, orange: t-test). Stars indicate the significance level (*: adjusted p<0.05, **: adjusted p<0.01, ***: adjusted p<0.001; Benjamini-Hochberg corrected), reference was set to Plasma, HMC chain length was set to 80,000 with a burn-in of 10,000.

### scCODA detects staining confirmed increase of disease-associated microglia in Alzheimer’s disease on few replicates

Second, we analyzed the compositional changes of three microglia cell types in an Alzheimer’s disease (AD) mouse model^15^ (**Fig. 2b**). Here, the number of replicates of sorted cells from cortex and cerebellum was low (n=2 per group), thus challenging standard statistical testing scenarios. In the cortex, scCODA identified statistically credible changes both in microglia 2 and disease-associated microglia (DAM) using the most abundant tissue-resident microglia 1 as reference cell type. By contrast, scCODA detected no statistically credible change in the cerebellum, which is known to be unperturbed in AD. Keren-Shaul et al.^15^ quantified the increase of DAM in the cortex of the AD mouse model via staining. While DAM localize in close proximity to amyloid-beta plaques and show a distinct inflammatory gene expression pattern, microglia 2 tend to represent an intermediate state between DAM and homeostatic microglia 1^15^ (**Supplementary Fig. 3**). Therefore, our analysis with scCODA supports the contribution of DAM in AD.

### scCODA scales to large sample sizes and cell type numbers

We next analyzed compositional changes of cell types in single-cell data from patients with ulcerative colitis (UC) compared to healthy donors^1^. Here, biopsy samples from the epithelium and the underlying lamina propria (**Fig. 2c**) were enzymatically separated and subsequently analyzed with scRNA-seq, resulting in 51 cell types from 133 samples. The epithelium and the lamina propria represent two different compartments and were tested separately. However, some epithelial cells ended up in the lamina propria samples and vice versa. For testing, we summarized these cells as non-epithelial in the epithelium and as epithelial in the lamina propria (**Fig. 2d-e**). We then re-analyzed the data with the Dirichlet regression model used in Smillie et al.^1^, leading to more statistically significant results compared to the original publication. Similar to the Dirichlet regression model, scCODA identified several statistically credible cell type changes in healthy tissue compared to both non-inflamed and inflamed tissue in both epithelium and lamina propria. In the epithelium, both Dirichlet regression and scCODA identified significant and statistically credible changes, respectively, in the absorptive and secretory lineage, but scCODA also identified an increase in enteroendocrine cells. In the lamina propria, B cell subpopulations showed several changes, e.g., a decrease of plasma B cells with disease (validated with stainings in Smillie et al.^1^), and an increase of follicular B cells. Moreover, consistent with our simulation studies demonstrating scCODAs higher sensitivity for lowly abundant cell types, scCODA uniquely detected statistically credible changes in several low-abundant immune cell populations. For instance, scCODA identified regulatory T cells (T_reg_) to be more abundant in UC patients which is consistent with other studies^17^. Smillie et al. combined the results of their Dirichlet regression with two non-compositional tests, Fisher exact test and Wilcoxon rank-sum test, to identify absolute changes in each population independently. Using such a two-stage procedure, Smillie et al. also reported changes in the low-abundant cell types such as T_reg_ cells. However, scCODA is more robust than Dirichlet regression in terms of MCC when we successively subsampled the number of donors from 29 (12 healthy donors and 17 UC patients, m=5 random subsamples) to 9 (2 healthy donors and 7 UC patients, m=5 random subsamples) in the epithelium test case (**Supplementary Fig. 4**), providing further evidence for scCODA’s ability to detect compositional changes in the small sample limit.

### scCODA detects cell type changes in COVID-19 patients that were not detected with non-compositional tests but confirmed in larger-scale studies

Next, we reanalyzed a recent COVID-19 single-cell study comparing compositional changes of major cell types in bronchoalveolar lavage fluid between healthy controls (n=4), severe (n=6) and moderate (n=3) COVID-19 cases^4^ using plasma cells as reference (**Fig. 2f**). The study originally reported significant differential changes in pDC’s between all groups, and depletion of T cells in severe cases vs. healthy controls using a t-test without multiple testing correction. scCODA confirmed these results but additionally identified a statistically credible depletion of T cells in severe vs. moderate cases, increases of NK and B cells in moderate vs. healthy cases, NK depletions in severe vs. moderate cases, as well as a decrease in mDCs in severe vs. moderate, and a Neutrophil increase in severe cases vs. moderate and healthy cases. Correlation of T cell abundances with severity is well established and has been used as risk factors for severe cases^18,19^. A decrease of NK cells with COVID-19 severity was observed between recovered and diseased patients^19^ and in PBMC through FACS analysis. Finally, higher Neutrophil proportions have been associated with severe outcomes^20^ and are suspected to be the main drivers of the exacerbated host response^21^, further confirming scCODA’s findings.

### scCODA accounts for the negative correlation structure for compositional changes and shows fewer false positives

Our final analysis considered a longitudinal scRNA-seq data set from the small intestinal epithelium in mice, studying the effects of *Salmonella* and *Heligmosomoides polygyrus* infection on cell type composition^6^. In contrast to the original Poisson regression data analysis^6^, scCODA found only a single statistically credible increase in Enterocytes in *Salmonella* infected mice **(Supplementary Fig. 5)**. The effects of *H. polygyrus* were also found to be less pronounced. While the Poisson model identified Tuft cells to be significantly increasing after three days of infection, scCODA only found statistically credible changes after ten days **(Supplementary Fig. 5)**. Both the Poisson model and scCODA identified Enterocytes as significantly and credibly affected, respectively. In addition, the Poisson model identified Goblet and early transit-amplifying cells as significantly changing which could not be confirmed by scCODA.

## Discussion

In summary, using a comprehensive set of synthetic and scRNA-derived compositional data sets and application scenarios, we established scCODA’s excellent performance for identifying statistically credible changes in cell type compositions. scCODA compared favorably to commonly used models for single-cell and microbiome compositional analysis, particularly when only a low number of experimental replicates are available or when detecting compositional changes with small effect sizes is of importance. We believe this is due to the Bayesian nature of the model as it adequately accounts for the uncertainty of observed cell counts, automatically performs model selection, and does not rely on asymptotic assumptions. scCODA not only correctly reproduced previously discovered and partially FACS-verified compositional changes in recent scRNA-seq studies, but also identified additional cell type shifts that were confirmed by independent studies, including T_reg_ cell enrichment in UC patients and Neutrophils increase in severe COVID-19 cases. Using synthetic benchmarks, we confirmed that standard univariate tests, such as Poisson regression models or t-tests, are inadequate for cell type analysis, since they do not account for the compositional nature of the data. While log-ratio transforms from compositional data analysis (such as the ALR used here) can partially mitigate these shortcomings, our Bayesian scCODA framework provided substantial performance improvements across all tested scenarios and is particularly preferable when only few replicates are available.

In its present form, our scCODA framework relies on pre-specified cell type definitions which, in turn, hinge on statistically sound and biologically meaningful clustering assignments. In situations where crisp clustering boundaries are elusive, for instance, due to the presence of the transient developmental processes underlying the data, joint modeling of different resolution hierarchies^22^ or modeling compositional processes^23,24^ may help account for such continuities changes. We believe that our scCODA framework offers an ideal starting point to model such advanced processes thanks to its hierarchical and extendable nature.

## Author contributions

M.B. and B.S. conceived of the study. J.O. developed scCODA and conducted the benchmarking study. M.B., J.O., and B.S. analyzed data. C.L.M. helped design the model comparison. B.S. and F.J.T. supervised the study and model development. B.S. designed the benchmarking study and conducted the power analysis. M.B., J.O., C.L.M., and B.S. wrote the manuscript. All authors read and approved the final manuscript.

## Acknowledgement

We would like to thank Malte Luecken for his support in the initial design of the study, as well as Karin Hrovatin and Lisa Sikkema for their support in developing synthetic data generation methods and testing the implementation of scCODA. F.J.T acknowledges financial support by the Bavarian Ministry of Science and the Arts in the framework of the Bavarian Research Association "ForInter" (Interaction of human brain cells). B.S acknowledges financial support by the Postdoctoral Fellowship Program of the Helmholtz Zentrum München.

## Online Methods

### Model description

We seek to identify the credibly associated covariates X^NxM^ to observed cell counts Y^NxK^ of K cell types measured in a single cell experiment with N samples and M covariates. We address this question with a Bayesian generalized linear multivariate regression framework using a Dirichlet Multinomial model with a log-link function to account for the compositional nature and uncertainties in the observed data. Covariate coefficients are hierarchically modeled using a non-centered parameterization^25,26^. For automatic model selection and identification of credibly associated covariates and affected cell types, we utilize a logit-normal prior as a continuous relaxation of the spike-and-slab prior^10^ resulting in the following hierarchical model:

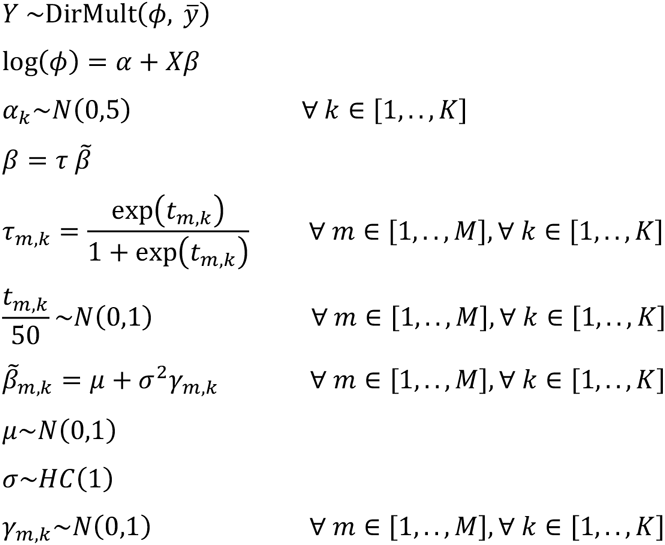

with *N* describing a Normal and *HC* a Half-Cauchy distribution following Polson et al.’s suggesting of hyperpriors for global scale parameters^27^.

To prevent identifiability issues of the covariate parameters, we reparametrize the model and choose one cell type as reference, forcing its covariates *β*_*k*_ = 0 as in Maier et al.^11^.

Parameter inference is performed via Hamiltonian Monte Carlo (HMC) using ten leapfrog steps per iteration with automatic step size adjustment according to Betancourt et al.^28^. Per default 20,000 iterations are performed with 5,000 iterations used as burn-in. The parameters *α*_*k*_, *γ*_*m,k*_ are randomly initiated by drawing from standard normal priors. *t*_*m,k*_ is always initialized with 0 to ensure unbiased model selection, while *μ* and σ^2^ are initialized with 0 and 1, respectively. If the data contains 0 entries, a pseudo-count of 0.5 is added to all cell counts to reduce numerical instabilities.

After parameter inference, we calculate the inclusion probability *P*_*inc*_(*β*_*k,m*_)of the covariates as follows:

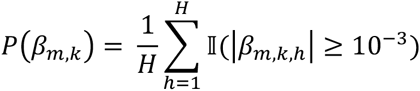

with H the number of HMC iterations and II the indicator function. To identify credibly associated covariates, we compare the calculated inclusion probabilities with a decision threshold *c*, which is by default set to *c* = 1 − 0.77 ⋅ *K*^−0.29^ **(Online Method - Spike-and-slab threshold determination)**.

### Simulation description

We carried out all benchmark studies by repeatedly generating compositional datasets 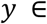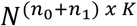 that have similar properties as the data from scRNA-seq experiments. For all synthetic datasets, we assumed a case-control setup with *n*_0_ and *n*_1_samples in the two groups and *K* cell types, as well as a constant number of cells ȳ in each sample.

We generated the synthetic datasets row-wise, with each row a sample of a Multinomial (MN) distribution *y*_*i*_ = *MN*(ȳ, *α*), and the probability vector *α* being a softmax of a multivariate normal (MVN) sample: *α* = *softmax*(MVN(*μ, Σ*)). We always used a covariance matrix of *Σ* = 0.05 *Id*_*k*_, which mimics the variances observed in the experimental data of Haber et al.^6^

The mean vector *μ* for each sample was calculated from the mean abundance of the first cell type in control samples (no effect) *μ*_0_, and the mean change in abundance of the first cell type between the two groups *μ*′. All other cell types were modeled to be equally abundant, leading to 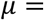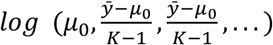 for control samples, and 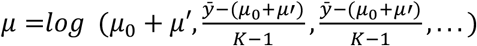 for samples in the other group.

For all three benchmark studies, we defined sets of values for all parameters mentioned above and generated *r* datasets for every possible parameter combination. We then applied scCODA with the last cell type chosen as reference to each synthetic dataset. For the model comparison benchmark **(Online Methods - Model comparison)**, all models were applied to all generated datasets.

The sets of generation parameters were as follows:

- Spike-and-slab threshold determination **(Supplementary Fig. 1a-e)**: {*K* = {2, 3,…, 15}; *n*_0_ = *n*_1_ = {1, 2,…, 10}(*only balanced setups – always n*_0_ = *n*_1_); {*ȳ* = 5000}; *μ*_0_ = 1/*K*; *μ*′ = {0.25, 0.5, 1} ∗ *μ*_0_; *r* = 20}
- Model comparison **(Fig. 1d)**: {*K* = 5; *n*_0_ = *n*_1_ = {1, 2,…, 10}(*only balanced setups – always n*_0_ = *n*_1_); {*ȳ* = 5000}; *μ*_0_ = {200, 400, 600, 800, 1000}; *μ*′ = {0.33, 0.5, 1, 2, 3} ∗ *μ*_i_; *r* = 20}
- Power analysis **(Fig. 1f)**: {*K* = 5; *n*_0_ = *n*_1_ = {1, 2,…, 10}(*also imbalanced setups*); {*ȳ* = 5000}; *μ*_0_ = {20, 30, 50, 75, 115, 180, 280, 430, 667, 1000}; *μ*′ = {10, 20, 30, 40, 50, 60, 70, 80, 90, 100, 200, 400, 600, 800, 1000}; *r* = 10}

### Spike-and-slab threshold determination

To identify statistically credible effects, scCODA compares the posterior inclusion probability

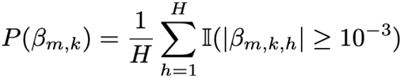

to a threshold *c*. To find a value for *c* that reliably identifies credible effects on single-cell population data, we applied scCODA to a range of 9,000 simulated data sets **(Simulation description)** and compared the inclusion probability for each cell type *k* to possible thresholding values ranging from 0 to 1. We measured the performance of scCODA by different binary classification metrics, noting a dependence on the number of cell types *K* in all metrics **(Supplementary Fig. 1a-d)**. To enable scCODA to evaluate datasets with dimensionality outside of the range of the simulation study, we determined the optimal thresholding values in terms of MCC for each *K* and approximated these as a function of the form *c* = 1 – *a* ⋅ *K*^*b*^. We found the optimal values for *a* and *b* by performing a grid search, using the sum of squared residuals as a metric **(Supplementary Fig. 1e)**. We excluded the case *K* = 2 from this optimization, as it represents an edge case where the only unchanged cell type is also the reference cell type. We found the optimal parameters to be *a* = 0.77, and *b* = −0.29.

### Power analysis

To be able to estimate the required sample sizes for an intended MCC, we fitted a linear regression model using sample size, absolute change in cell count, log-fold change, and all pairwise interactions using the simulation results **(Online Methods - Simulation description)**. To this end, we min-max and empirical logit transformed^29^ the measured performance to translate the MCC values to an unconstrained domain:

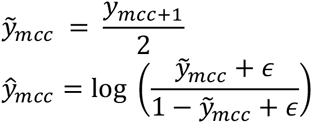

with *∊* being the minimal min-max transformed MCC reached during the benchmark simulation.

We performed a backward model selection with repeated 10-fold cross-validation to reduce the feature set. The final model consisted of sample size, log fold-change, absolute cell count change, and the interaction effect between absolute cell count and log-fold change, which was expected given that we observed an interaction effect between these two variables in the raw benchmark results (**Supplementary Fig. 1d-f**).

With the fitted model, we can now inverse estimate the required total sample size *x*_*ss*_ with fixed MCC *ŷ*_*mcc*_, log fold-change *x*_*fc*_, and absolute cell count change *x*_*cc*_ as:

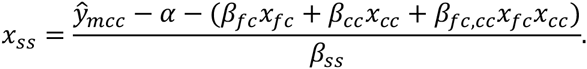

With this formula, we estimated the sample size for a fixed MCC of 0.8 across changing log fold-changes between [0, 5] and the fraction of cell type sizes to total cell counts between [0.01, 0.2].

### Analysis of publicly available data sets

#### Single-cell RNA-seq data of PBMCs from supercentenarians

We downloaded the processed single-cell RNA-seq count matrices comprising PBMCs of seven supercentenarians and five younger controls from http://gerg.gsc.riken.jp/SC2018/. Read counts were log-transformed and PCA embedded using the first 50 PCs. Leiden clustering was used to cluster cells into major groups. Following the described analysis in Hashimoto et al.^3^, we annotated the major cell types including T cells characterized by *CD3* and T cell receptor (*TRAC*) expression, B cells characterized by *MS4A1* (*CD20*) and *CD19* expression, natural killer cells characterized by *KLRF1* expression, monocytes characterized by *CD14* and *FCGR3A* (*CD16*) expression, respectively, and Erythroblasts characterized by *HBA1* expression, and determined their cell counts per sample (**Supplementary Fig. 2**). All analysis steps were carried out using scanpy v.1.5.1.

#### Single-cell RNA-seq data of microglia in Alzheimer’s disease (AD) mouse model

We downloaded the raw single-cell RNA-seq count matrices (deposited at GEO, accession code GSE98969) comprising immune cells isolated from the mouse brain in wild type (WT) and AD mice^15^. The complete dataset with all samples consists of 37,248 cells. We filtered out ERCC spike-ins before computing the quality metrics of all cells. We then excluded 12,053 cells with less than 500 UMI counts and 11,065 genes, which were not expressed. We subsequently normalized by library size with target sum 10,000 counts (CPM normalization) and log+1 scaled. Following the analysis of Keren-Shaul et al.^15^, we selected the samples of six-month-old mice from AD and WT, which haven’t been sorted by brain region, resulting in 9,196 cells. It must be noted that Keren-Shaul et al. reported 8,016 cells when they first annotated immune cells in six-month-old mice (see Figure 1 in Keren-Shaul et al.). We evaluated batch effects based on the clustering results and visual inspection of the UMAP plots, where none of the samples clustered separately in any of the clusters, which is, in this case, sufficient to obtain cell types. We clustered the data using Louvain clustering with resolution 1 and annotated cell types using the previously reported marker genes as microglia 1 (*CTSD, CD9, HEXB, CST3*), microglia 2-3 (*LPL, CST*), granulocytes (*CAMP, S100a9*), T/NK cells (*S100a4, NKG7, Trbc2*), B cells (*RAG1, CD79b, CD74*), monocytes (*S100a4, CD74*), perivascular macrophages (*CD74, CD163, MRC1*) (see **Supplementary Fig. 3**). We subsequently sub-clustered the microglia population into three clusters, assigning the labels microglia 1, 2, and 3, respectively. Similar to Keren-Shaul et al, we assigned the region-sorted samples of AD and WT mouse model (n=2 per region) with a k-nearest neighbor classifier (k=30). We then evaluated the number of unassigned cells, performed another round of Louvain clustering and assigned the remaining cells based on majority vote for the clustering result, i.e., when unassigned cells clustered predominantly with microglia 1, they were all assigned to microglia 1. The obtained proportions of microglia subpopulations are in accordance with the previously reported proportions. All analysis steps were carried out using scanpy v.1.5.1.

#### Single-cell RNA-seq data of ulcerative colitis in human donors

We used the annotated single-cell RNA-seq data of the colon epithelium from 12 healthy donors and 18 patients with chronic inflammation^1^. From healthy donors, samples from 2 adjacent locations were taken. From patients, biopsies from inflamed and adjacent normal tissue (‘non-inflamed’) were taken. Further, the biopsies were separated by enzymatic digestion into the epithelium (‘Epi’) and the lamina propria (‘LP’) before single-cell RNA-sequencing. The study comprises a total of 365,492 transcriptomes from 133 samples. The data were downloaded from Single Cell Portal (accession ID SCP259, https://singlecell.broadinstitute.org/single_cell/study/SCP259). The analysis code and description were provided at https://github.com/cssmillie/ulcerative_colitis.

The original study annotated all cell types together, resulting in 51 different cell types. However, some cell types that are originally located in the LP have been found in the epithelial samples and vice versa. For the differential composition analysis of the Epi and LP, we considered the non-epithelial and epithelial cell types, respectively, as one group. Therefore, we tested the changes in 16 cell types in the Epi and 37 cell types in LP. In addition, we re-analyzed the data using the Dirichlet regression model as in Smillie et al.^1^ (with R package DirichletReg v.0.7-0 in R v.3.5.2). Importantly, we realized that Smillie et al. summed up the counts of the same replicates (as described in the analysis scripts in https://github.com/cssmillie/ulcerative_colitis), while we consider every replicate as independent. Overall, we have data from 29 donors (61 samples, where 24 healthy, 21 non-inflamed, 16 inflamed) in Epi and data from 30 donors (72 samples, where 24 each healthy, non-inflamed and inflamed, respectively) in LP. Furthermore, we tested the impact of adding a pseudocount of 0.5 to the data to increase the acceptance rate in scCODA and the chain length in the HMC sampling (**Supplementary Fig. 6**). Specifically, we tested chain lengths of 20,000, 40,000, 80,000, and 150,000 iterations with a burn-in of 10,000 in the Epi case, while we tested chain lengths of 200,000, 400,000, and 800,000 iterations with a burn-in of 10,000 in the LP case due to the larger number of cell types in LP compared to Epi.

For the subsampling analysis in scCODA and Dirichlet regression, we used the Epi data and excluded 12 to 20 donors with all corresponding replicates. The dataset contained 29 donors (12 healthy, 17 UC patients), and we successively reduced the number of donors in the dataset from 17 to 9, where one donor from each group was removed in each step (stratified subsampling). Hence, when the subsampled dataset with 17 donors consisted of 6 healthy donors and 11 UC patients, while the dataset with 9 donors consisted of 3 healthy donors and 6 UC patients. We repeated each round 5 times with a different subset of donors to reduce selection bias (**Supplementary Fig. 4**).

#### Single-cell RNA-seq data of bronchoalveolar immune cells in patients with COVID-19

We used the annotated single-cell RNA-seq data of the bronchoalveolar lavage fluid cells from three patients with moderate COVID-19 progression, six patients with severe COVID-19 progression, four healthy donors, and a publicly available sample^4^. The cell type annotations of all samples were provided at https://github.com/zhangzlab/covid_balf.

#### Single-cell RNA-seq data of small intestinal epithelium cells infected with different bacteria

Annotated single cell transcriptomics data of epithelial cells from the small intestine of mice infected with three different bacterial conditions were downloaded from Single Cell Portal (accession ID SCP44, https://singlecell.broadinstitute.org/single_cell/study/SCP44). The data consisted of a control group of four mice (3,240 cells total) and three groups of two mice each, measured after two days for *Salmonella* (1,770 cells total), as well as three (2,121 cells total) and ten days (2,711 cells total) after *H. polygyrus* infection, respectively.

### Model comparison

We compared scCODA’s ability to correctly identify significant compositional changes in a setting typical for single-cell experiments to other methods recently used in scRNA-seq analysis and approaches from the field of microbial population analysis. We applied all methods to each of the 5,000 datasets generated for the comparison analysis **(Online Methods - Simulation description)** and recorded which of the 5 components each method found to statistically differ between the two groups. We then compared these results to the ground truth assumption of only the first component changing significantly via binary classification metrics (significant vs. non-significant changes). We chose Matthews’ correlation coefficient as our primary metric, as it best accounts for the numerical imbalance between the two groups. Details on the individual methods can be found in **Supplementary Table 1**.

### Implementation

The method has been implemented in Python 3 using Tensorflow >= 2.3.1^30^, Tensorflow-Probability >= 0.11^31^, ArviZ >= 0.9^32^, numpy >= 1.19, and Scanpy >= 1.5^33^. The Power Analysis was performed using Scikit-learn >= 0.23^34^ (Python 3.8) and caret package^35^ (R 3.7).

Source code can be found at https://github.com/theislab/sccoda. All code to reproduce the presented analyses can be found at https://github.com/theislab/scCODA_reproducibility. Simulated data and benchmark analyses results are available at https://doi.org/10.5281/zenodo.4305907

---

**Supplementary Figure 1.**
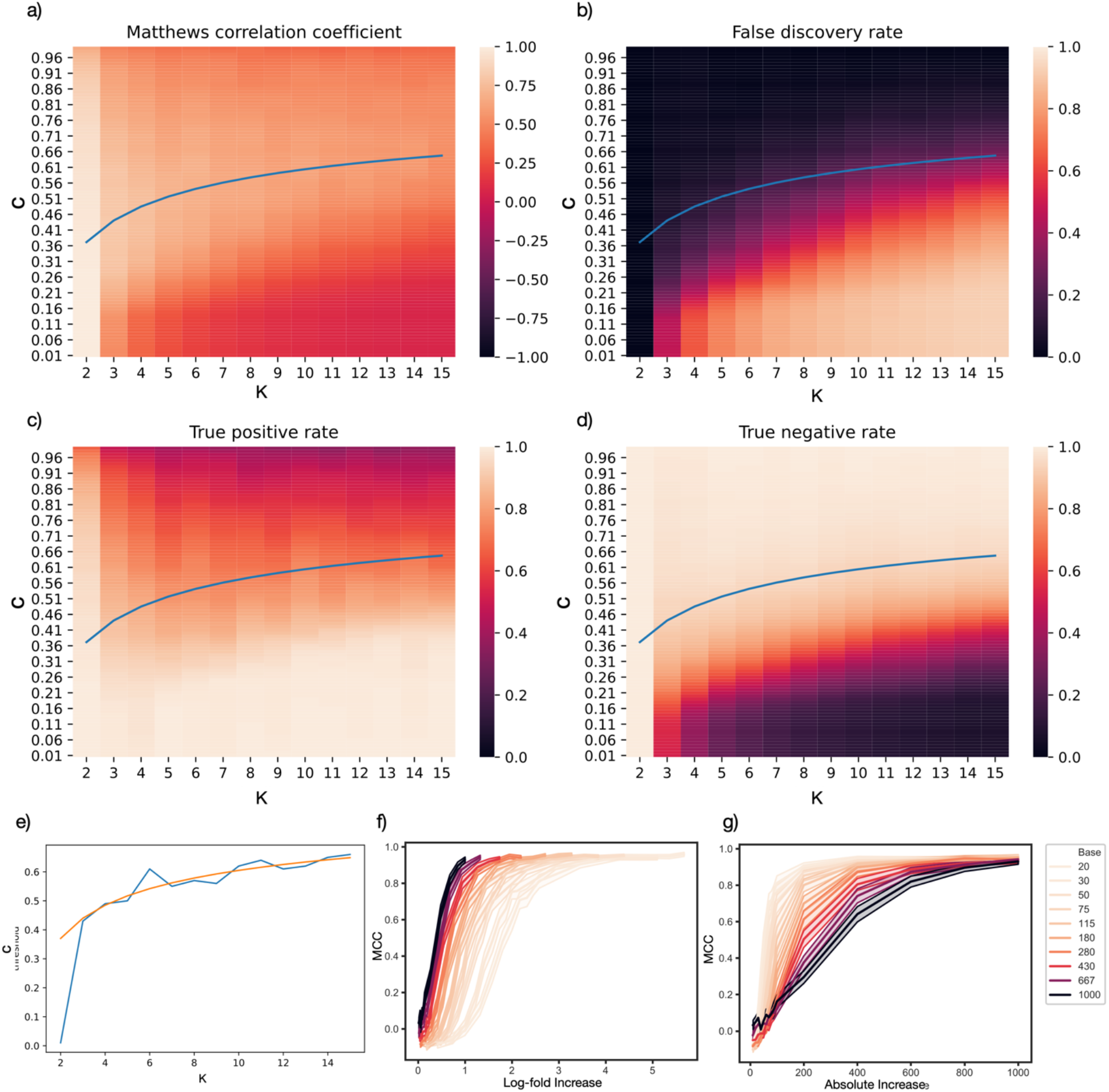
Benchmark evaluation results for spike-and-slab threshold determination **(a-e)** and overall benchmark **(f-g) (Online Methods - Simulation Description)**. The optimal thresholding value for the posterior inclusion probability of the spike-and-slab prior on each effect depends on the number of cell types. The blue line always represents the chosen optimal thresholding function *c* = 1 – 0.77 ⋅ *K*^−0.29^ **(Online Methods - Spike-and-slab threshold determination)**. Benchmark performance for different numbers of cell types and thresholding values in terms of **(a)** Matthews correlation coefficient, **(b)** false discovery rate, **(c)** true positive rate, and **(d)** true negative rate. **(e)** The optimal threshold function (orange) closely matches the MCC-maximizing threshold values (blue) for all numbers of cell types. **(f)** The performance of scCODA (measured by MCC) depends on the amount of change in abundance. The “Base” value represents the mean cell count of the only differentially abundant cell type. For cell types with higher initial abundance, scCODA can detect smaller relative (log2-fold) changes between the two groups. **(g)** For cell types with higher initial abundance, the absolute (count) change must be higher to reliably detect changes in abundance.

**Supplementary Figure 2:**
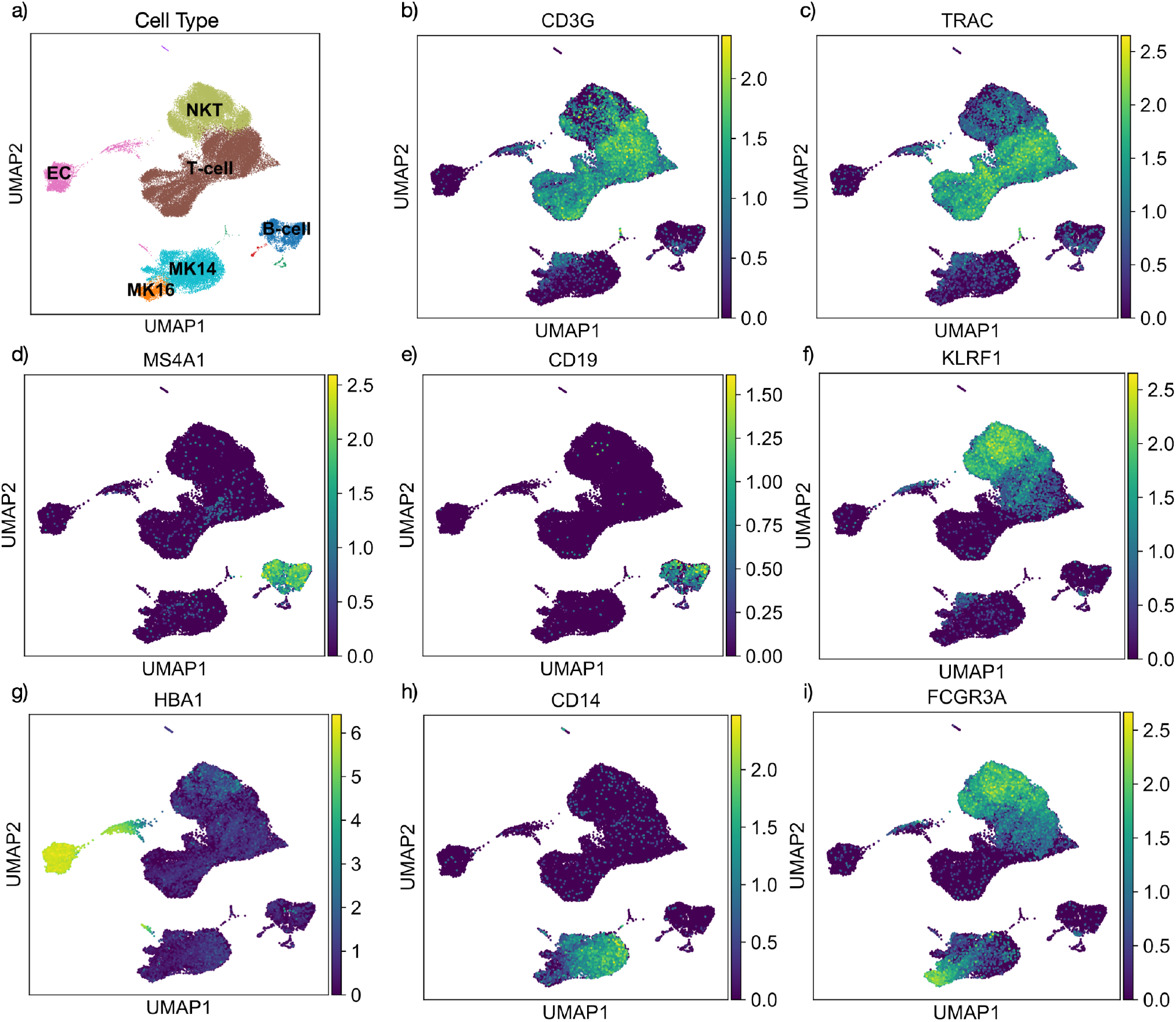
Re-analysis of supercentenarian data of Hashimoto et al.^3^. **(a)** Final annotation of major cell types. **(b-i)** expression pattern of *CDR3* and *TRAC* identifying T-cells, *MS4A1* and *CD19* identifying B-cells, KLRF*1* natural killer cells (NKT), *HBA1* Erythroblasts (EC), *CD14* and *FCGR3A* (CD16) Monocyte subtypes (CD14+, CD16+, denoted as MK14 and MK16).

**Supplementary Fig. 3:**
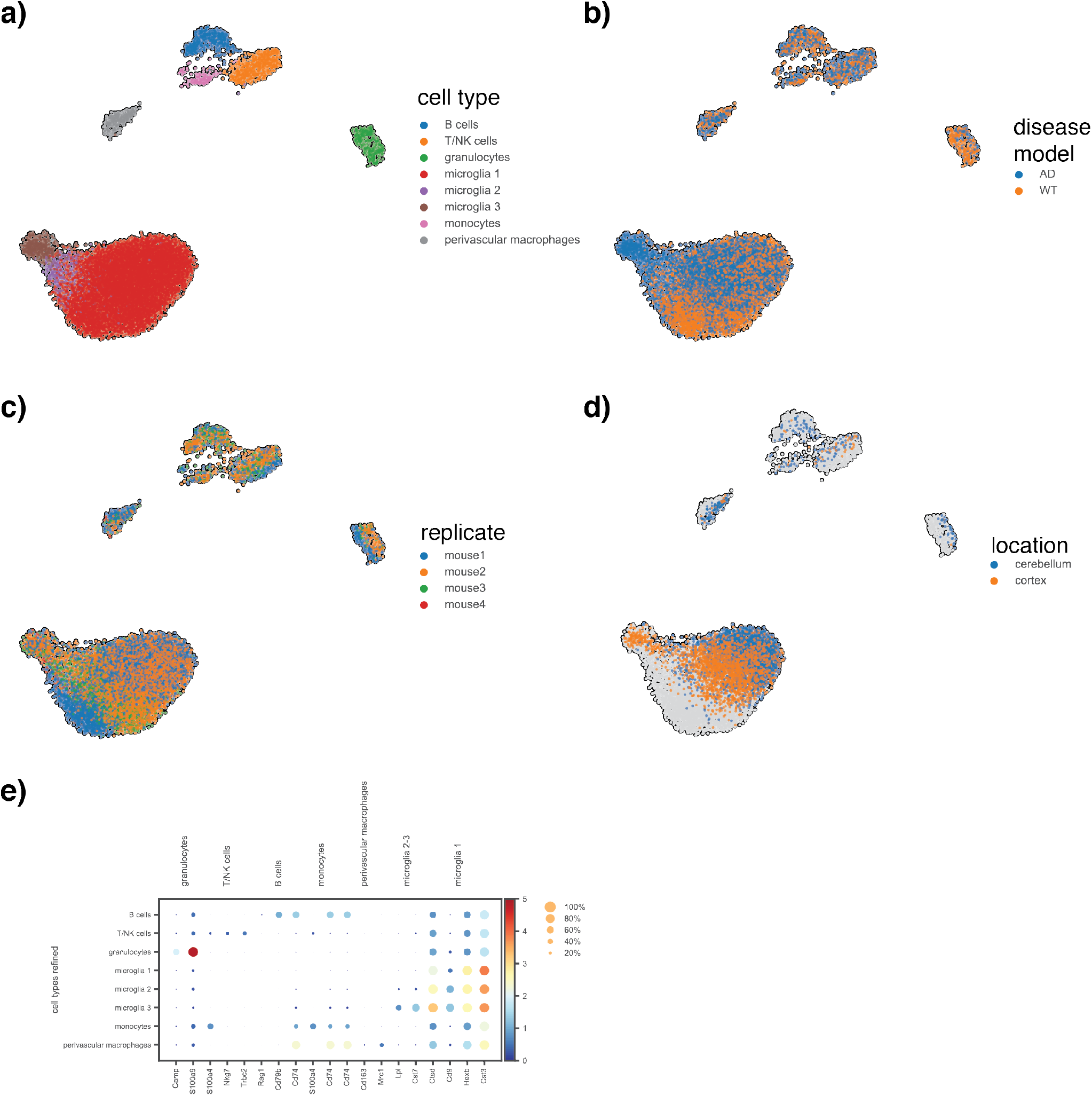
Re-analysis of microglia data in Alzheimer’s disease (AD) mouse model^15^. **(a)** Joint cell type annotation of cells. **(b)** Cell distribution in the both wild type (WT) and AD mouse model. **(c)** Distribution of cells from different replicates does not indicate strong batch effects. **(d)** Location of cells sorted from cortex and cerebellum. Location of grey cells was not reported. **(e)** Dot plot of marker gene expression of the annotated cell populations **(a)**.

**Supplementary Figure 4:**
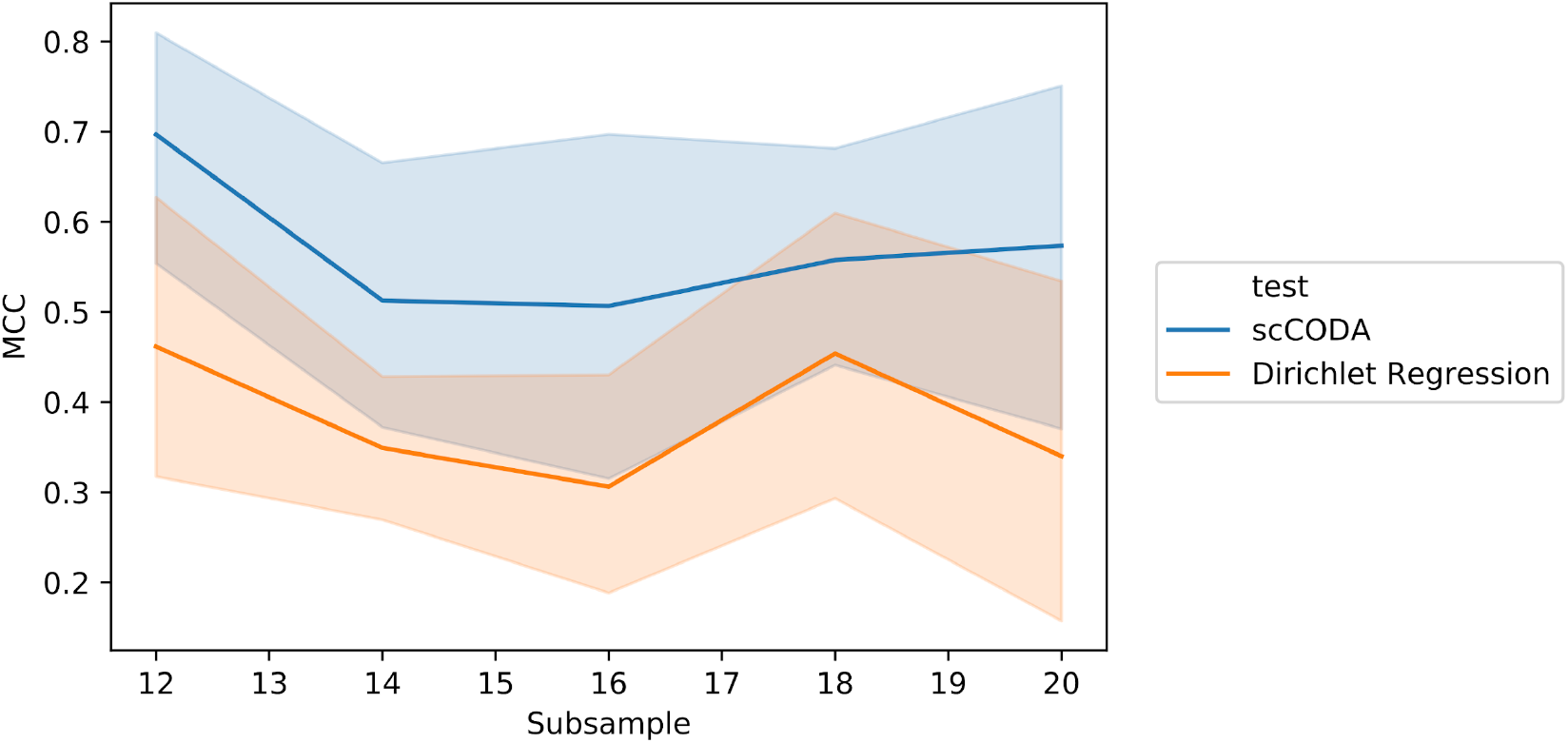
Robustness of the results of scCODA and Dirichlet regression upon subsampling of donors in epithelium data of Smillie et al. The dataset contained 29 donors (12 healthy, 17 UC patients), and we successively reduced the number of donors in the dataset from 17 to 9 (corresponds to subsample 20 in the plot), where one donor from each group was removed in each step. Each subsampling step was repeated five times. Ground truth for MCC computation is the set of statistically differentially abundant cell types in the full dataset.

**Supplementary Figure 5:**
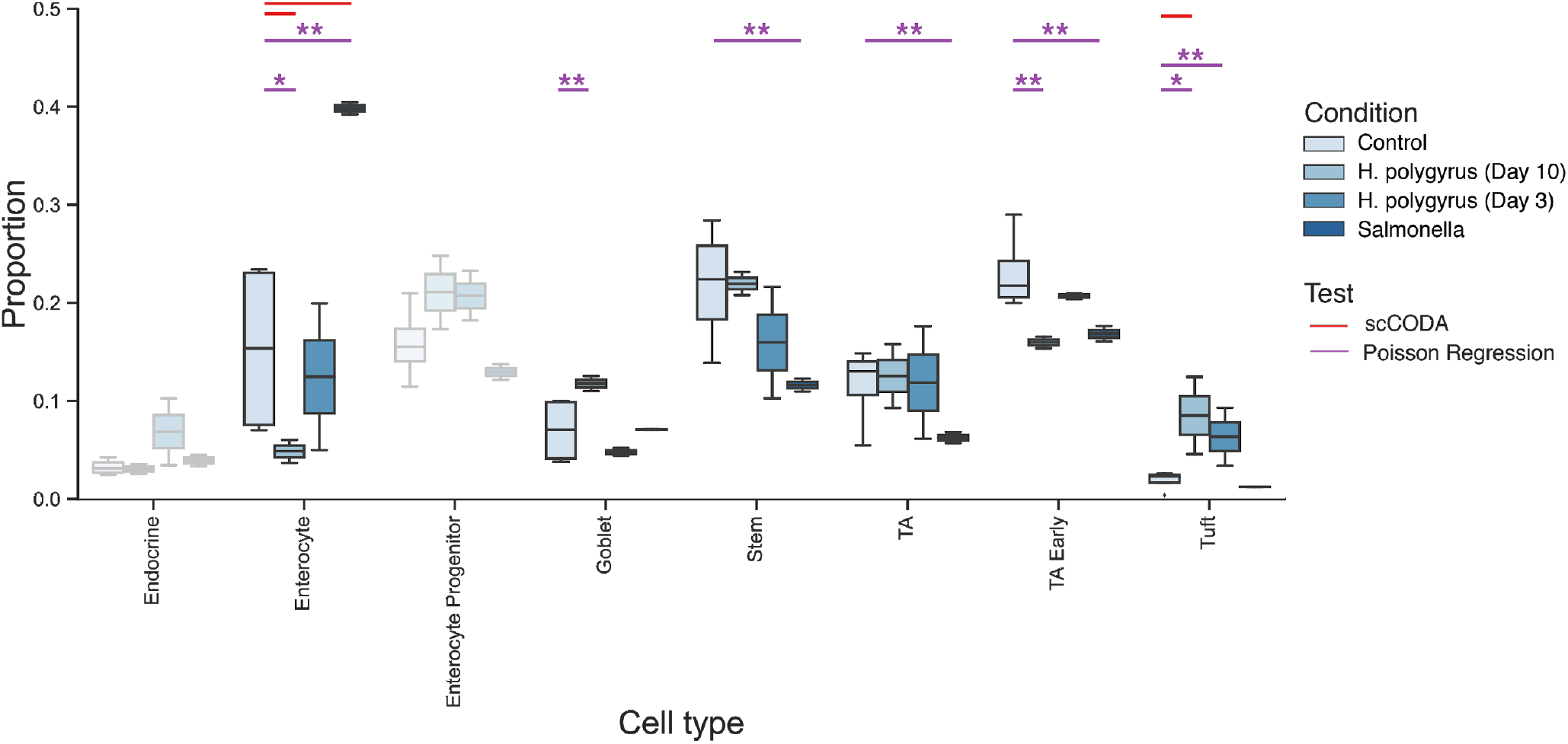
Compositional analysis of Haber et al.^6^ on the response to pathogen infection in the small intestinal epithelium of the mouse. Significant and credible results are depicted as colored bars (Red: scCODA, purple: Dirichlet regression), stars depict the significance of the Dirichlet regression model (*: p<10^−5^, **: p<10^−10^).

**Supplementary Figure 6:**
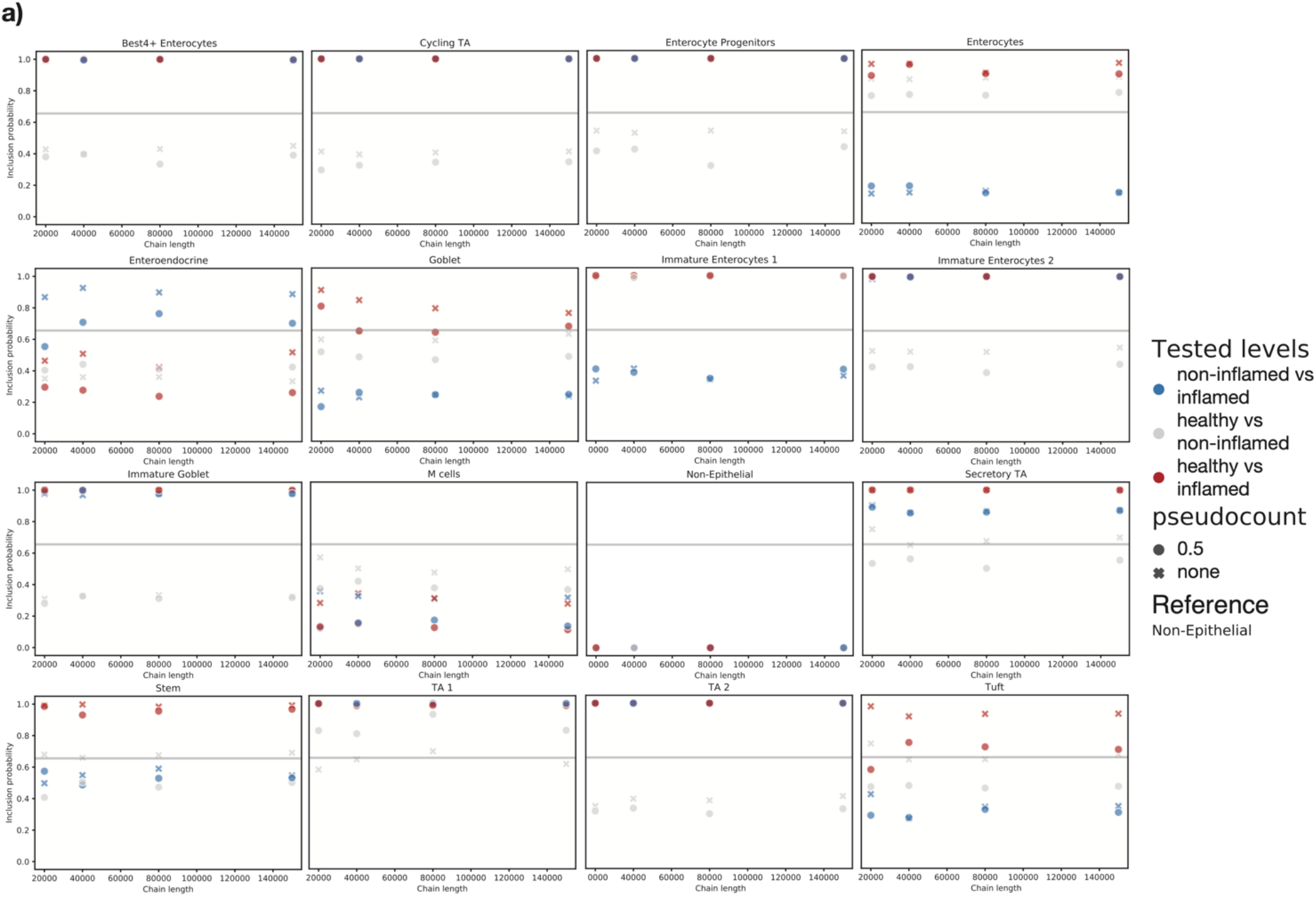
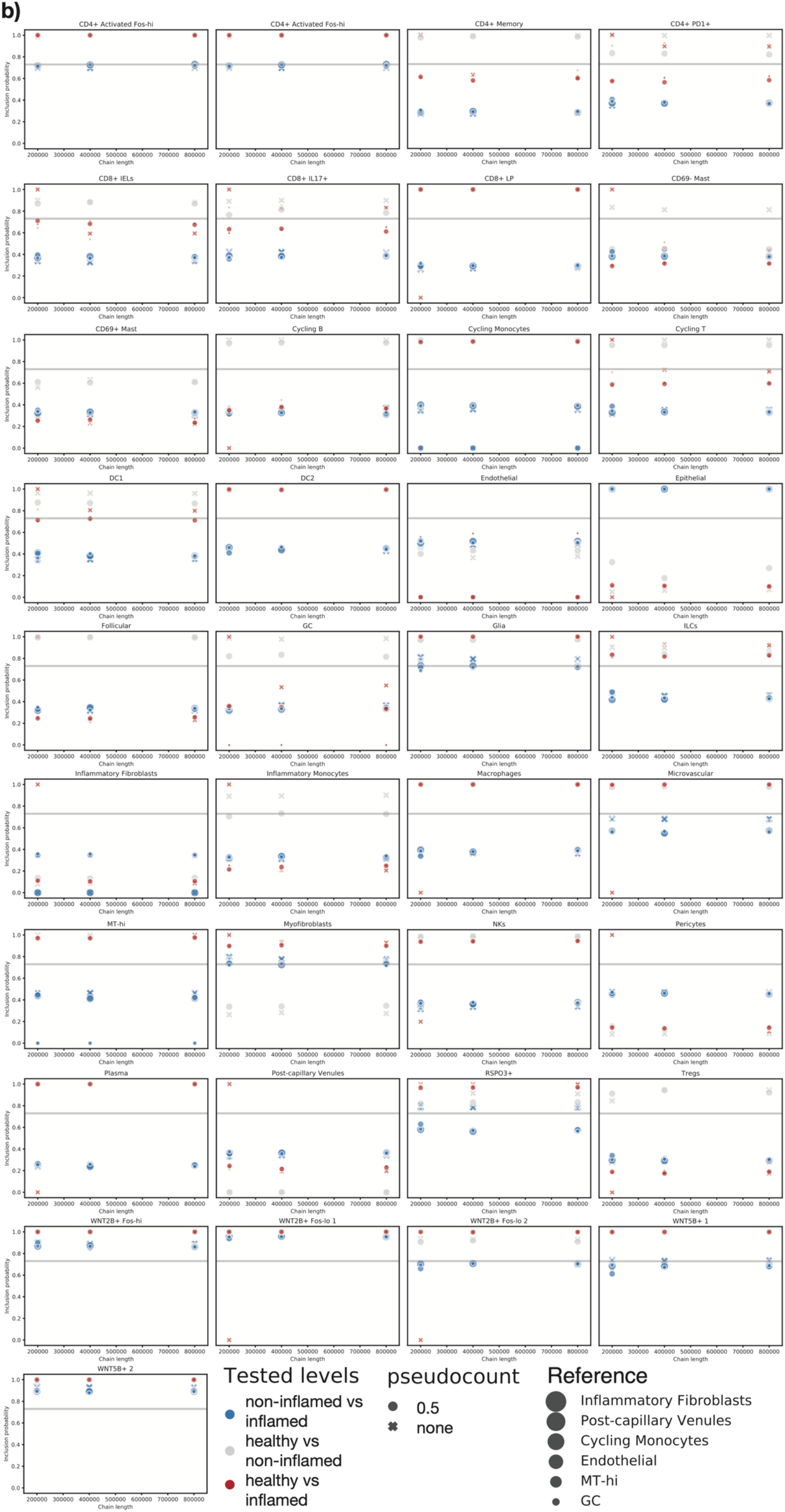
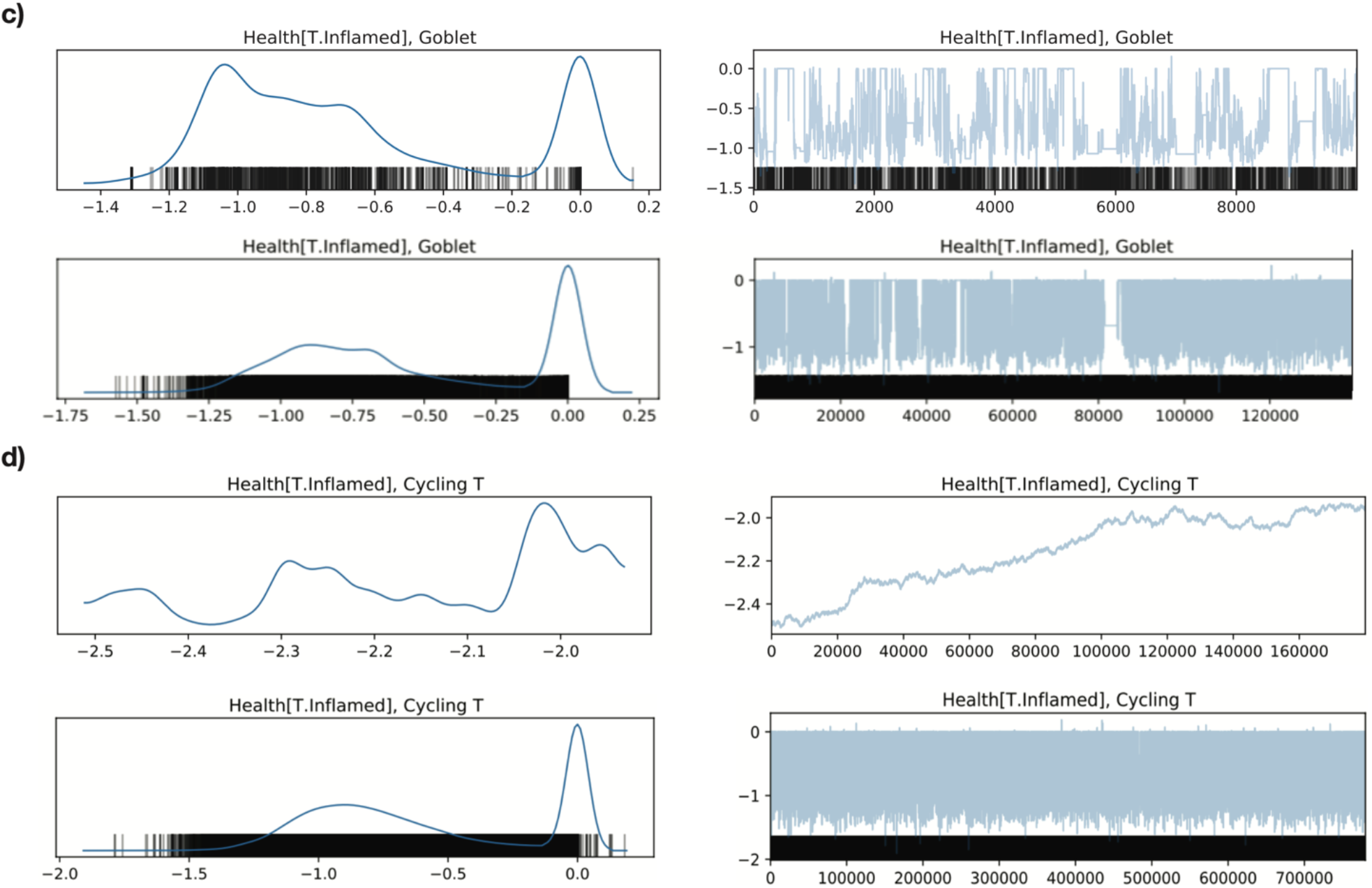
Convergence of HMC sampling for many cell types (in data of Smillie et al.). **(a-b)** Inclusion probabilities for pairwise tests in the epithelium **(a)** and lamina propria **(b)** of healthy donors and patients of UC. Colors depict the tested levels, symbols depict the addition of pseudocounts, symbol sizes depict different references. The effect of the reference is set to zero. **(c-d)** Trace plots of different chain lengths for the parameter inference in Goblet cells with pseudocount comparing healthy and inflamed samples (reference Non-Epithelial) **(c)** and Cycling T cells without pseudocount comparing healthy and inflamed samples (reference Endothelial) **(d)**.

**Supplementary Table 1.** Methods and configurations used in the benchmark comparison.

| Name | Description | Implementation details |
| --- | --- | --- |
| scCODA | Our proposed method | Reference cell type always set to the last component |
| Standard Dirichlet-Multinomial | Fully Bayesian model: Log-linear model on components of a Dirichlet-Multinomial distribution. Selection of a reference cell type. HMC inference setup identical to scCODA. | Reference cell type always set to the last component |
| scDC | Single-cell differential composition analysis <sup>7</sup><br>*Note: This method did not give results for all datasets. The erroneous results were left out of the analysis.* | Number of bootstrap samples generated for each data set: 100; Significance level: 5% |
| ancom | Analysis of composition of microbiomes <sup>12</sup> | Default settings |
| ALDEx2 | ANOVA-Like Differential Expression tool for high throughput sequencing data <sup>36</sup>. Reference cell type set to the last component instead of the geometric mean; testing via t-test (Benjamini-Hochberg-corrected) | False discovery rate: 5% |
| ALR+t-test | Additive log-ratio transform of data, t-test on all components, Benjamini-Hochberg correction of p-values | Reference component: Last cell type; False discovery rate: 5% |
| ALR+Wilcoxon | Additive log-ratio transform of data, Wilcoxon-rank-sum test on all components, Benjamini-Hochberg correction of p-values | Reference component: Last cell type; False discovery rate: 5% |
| Dirichlet regression | R-Package DirichletReg <sup>11</sup> | Default settings; Significance level |
| t-test | t-test on all components of untransformed data, Benjamini-Hochberg correction of p-values | False discovery rate: 5% |
| Poisson regression | Poisson regression model used by Haber et al. <sup>6</sup>, Benjamini-Hochberg correction of p-values | False discovery rate: 5% |

